# Cytogeography of *Gagea bohemica* (Liliaceae) outside the Mediterranean: two ploidy levels, spatial differentiation of cytotypes, and occurrence of mixed-ploidy populations

**DOI:** 10.1101/2023.03.29.534790

**Authors:** David Horák, Bohumil Trávníček, Gergely Király, Jacqueline Détraz-Méroz, Tomáš Vymyslický, Marianthi Kozoni, Dörte Harpke, Michal Hroneš

**Affiliations:** Department of Botany, Faculty of Science, Palacký University in Olomouc, Šlechtitelů 27, CZ-78371 Olomouc, Czech Republic; Faculty of Forestry, University of Sopron, Bajcsy-Zs. u. 4., H-9400 Sopron, Hungary; Route de la Biolette 8, CH-1996 Basse-Nendaz, Switzerland; Agricultural Research Ltd., Zahradní 1, CZ-664 41 Troubsko, Czech Republic; Department of Agriculture, Faculty of Agriculture Forestry and Natural Environment, Aristotle University of Thessaloniki, GR-54124 Thessaloniki, Greece; Leibniz Institute of Plant Genetics and Crop Plant Research (IPK), Corrensstraße 3, D-06466 Seeland-Gatersleben, Germany

**Keywords:** absolute genome size, chromosome number, cytotype, flow cytometry, geophyte, pollen stainability, polyploidy

## Abstract

*Gagea bohemica* s. lat. is a morphologically and karyologically highly variable group with many morphologically similar “narrow” taxa currently considered as a single variable species. It is predominantly distributed in Mediterranean and warmer parts of temperate belt of Europe. The large-scale data on its cytogeography and population cytotype structure which could provide a basis for taxonomy is lacking, only scattered data on ploidy have been published from various parts of its range. In this study, we sampled 106 populations in broader Central Europe, the northeastern Balkan Peninsula and the northwestern Black Sea coast in order to analyse their ploidy level, genome size and pollen stainability. Two cytotypes, i.e., tetraploid (2n = 48) and pentaploid (2n = 60), were found in the study area using chromosome counting and flow cytometry, both in pure and mixed-ploidy populations. Pure pentaploid populations are mainly distributed in Austria, Czechia, northwestern Hungary and Slovakia while tetraploid cytotype in pure and mixed-ploidy populations forming two lineages which are concentrated into two disjunct geographical areas: a western lineage in Germany and Switzerland, and an eastern one in Bulgaria, southeastern Hungary, northern Greece, Romania, Serbia and Ukraine. The two lineages differ in their genome size regardless of their ploidy, indicating their independent origin. Analysis of pollen stainability using a modified Alexander stain revealed an unusual pattern with tetraploids having a lower pollen stainability (mean 44.29 %) than pentaploids (mean 70.70 %) but the western and eastern populations differed again from each other.

## Introduction

Polyploidy (a state with more than two chromosome sets in the nucleus) is an important diversifying mechanism in angiosperms that is significantly linked to the genetic diversity, phenotypic variation, ecology and reproduction strategies of particular species (e.g., Soltis 2005; Hull-Sanders et al. 2009; Soltis et al. 2009; Wood et al. 2009; Sonnleiter et al. 2016; Van Drunen and Husband 2019). Species with intraspecific ploidy variation (i.e., mixed-ploidy species, Kolář et al. 2017) provide insight into polyploid evolution within the frame of a recently shared history and genetic background. One of the consequences of polyploidy may be a niche shift which may be manifested by a different distribution and/or ecology of cytotypes (e. g., Karunarathne et al. 2018; Castro et al. 2020; Duchoslav et al. 2020; Urfus et al. 2021).

Although the frequency of polyploidy in Liliaceae is rather low compared to other monocot bulbous families (e.g., Asparagaceae and Amaryllidaceae), there are several genera with a high proportion of polyploids, i.e., *Clintonia* Raf., *Amana* Honda and especially *Gagea* Salisb. (Peruzzi et al. 2009). In the latter genus, the extent of polyploidy is very high with exclusively diploid species in early diverging groups up to undecaploids (11x) in terminal groups (Peruzzi 2012). The existence of entirely polyploid terminal groups and various polyploid series in several species illustrates the dynamic genome evolution in *Gagea*. The karyological data of *Gagea* were reviewed by Peruzzi (2003, 2008). Later, flow cytometry was used to estimate the genome size and infer the ploidy level of several species by Zarrei et al. (2012), Peruzzi et al. (2015) and Zonneveld et al. (2015). But more detailed information on cytogeography which may provide insights into the cytotype population structure at a larger geographical scale for *Gagea* species is still very scarce.

*Gagea bohemica* s. lat. (*Gagea* sect. *Didymobulbos* (K.Koch) Boiss.) is a bulbous geophyte with two basal leaves, alternate cauline leaves which are spread more or less evenly along stem and glabrous to hairy pedicels (Fig. 1A–D; Richardson 1980; Slater 1990; Hrouda 2010). It is widespread but disjunctively distributed in central and western Europe and the wider northeastern regions of the Mediterranean and Black Sea. It may also occur in North Africa, but the current presence of populations and their taxonomic status are unclear (Slater 1990; Peterson et al. 2010a; Fig. 2A). The basic chromosome number of *G. bohemica* s. lat. is assumed to be x = 12 (Peruzzi 2012) as in the entire tribe Tulipeae of Liliaceae. Based on various reports, it forms a diploid-polyploid complex with five ploidy levels ranging from diploid to hexaploid (Peruzzi 2003, 2008; Fig. 2A). According to published chromosome counts, cytotypes with lower ploidy levels are predominantly distributed in the Eastern Mediterranean, the Iberian Peninsula and France and Corsica, while the Apennine Peninsula and more northern areas of Europe are mainly occupied by cytotypes with higher ploidy levels (Peruzzi 2003, 2008; Gutiérrez and Vázquez 2010). However, information on the ploidy level of *G. bohemica* s. lat. outside the Mediterranean basin is rather scarce or completely absent in some parts of its range. Given that different ploidy levels can lead to different phenotypes and affect the reproductive strategy of *G. bohemica* s. lat. (Hrouda 2010; Jakab and Molnár 2011), it is important to know the distribution of cytotypes and their composition in populations.

**Fig. 1.**
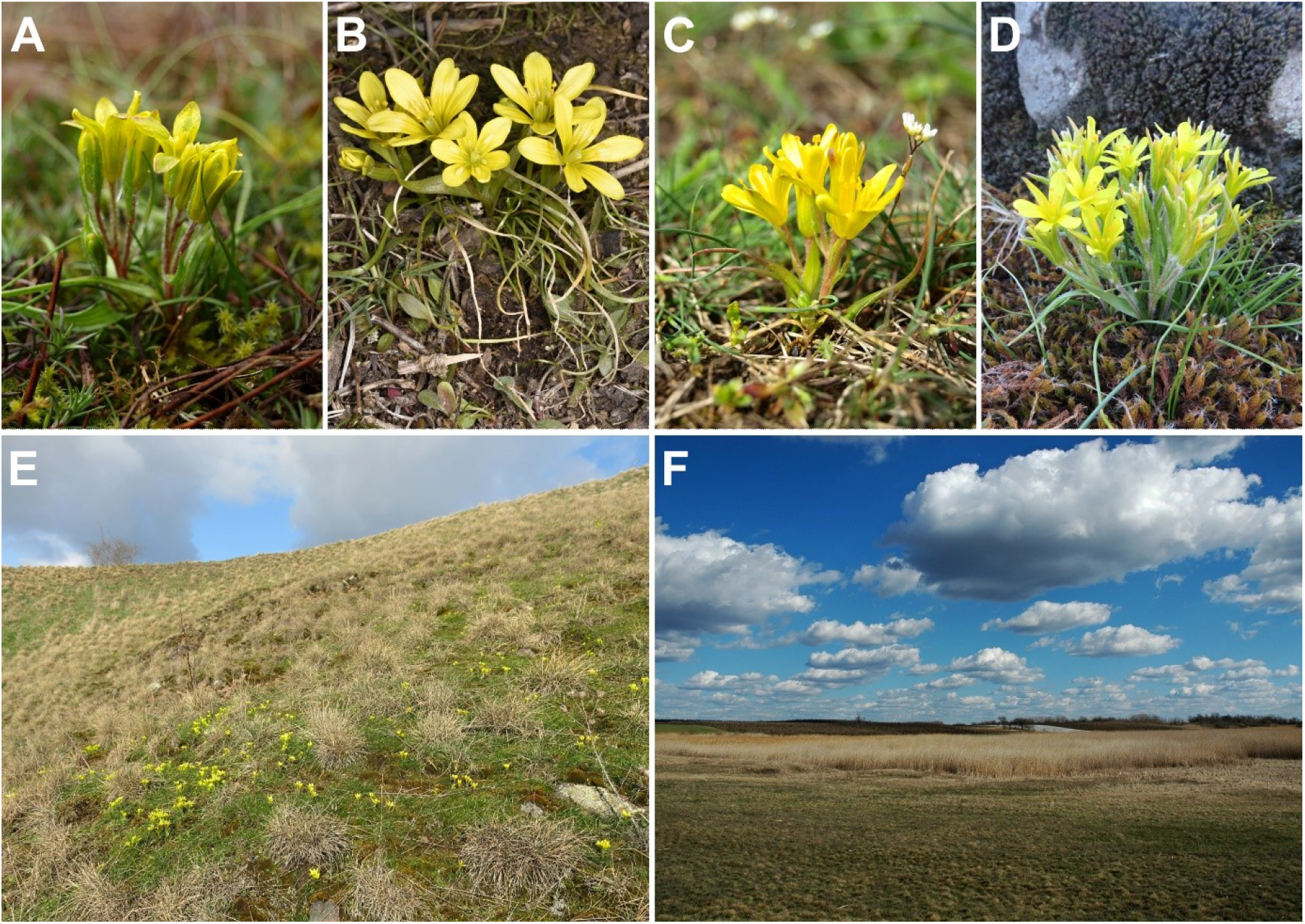
Variation in general habit of studied populations of *Gagea bohemica* and examples of their habitats. A – Germany, Neu-Bamberg (BOH45). B – Czech Republic, Radim (BOH43). C – Hungary, Kunszentmárton (BOH21). D – Bulgaria, Karnobat (BOH87). E – Germany, Mücheln (BOH64). F – Hungary, Kajánújfalu (BOH20). Authors of photographs: M. Hroneš (A–C), D. Horák (D), B. Trávníček (E) and T. Vymyslický (F).

**Fig. 2.**
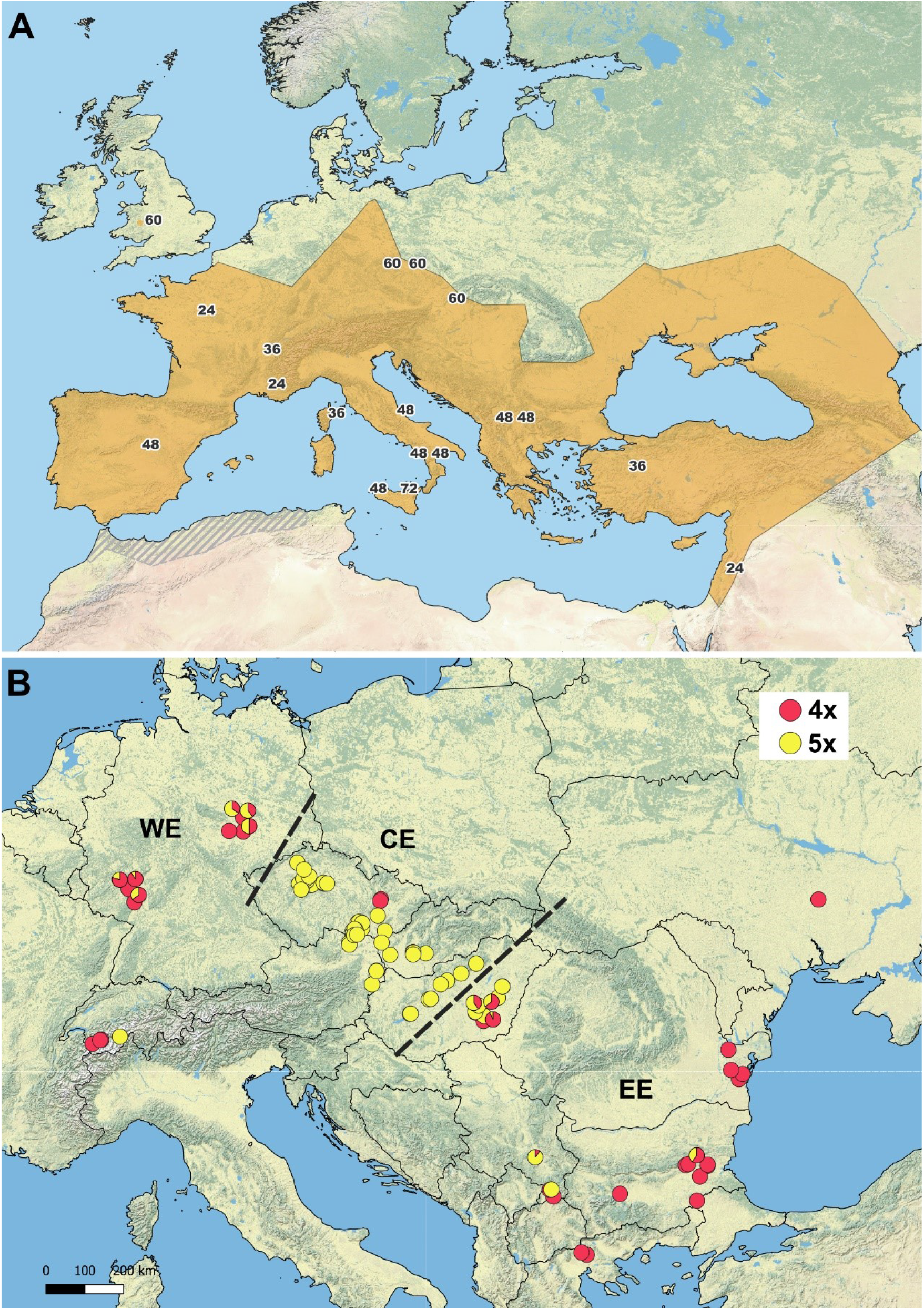
A – Approximate distribution of *Gagea bohemica* s. lat. (orange area; doubtful distribution in northern Africa marked by grey stripes) with marked locations of published chromosome counts; numbers indicate the published chromosome number (after Contandriopoulos 1962, Měsíček and Hrouda 1974, Heyn and Dafni 1977, Murín and Májovský 1983, Guerlesquin 1985, Sopova et al. 1984a, b, Slater 1990, Tison 1996, Peruzzi 2008, Gutiérrez and Vázquez 2010). B – Distribution of cytotypes of *Gagea bohemica* outside the Mediterranean found in this study (red – tetraploid cytotype; yellow – pentaploid cytotype; pie charts – mixed-ploidy population with a proportion of each cytotype; CE, EE, WE, striped line – division into three population groups further discussed in the text).

Taxonomy of *G. bohemica* s. lat. is a matter of ongoing debate. Taxonomic concepts ranging from recognition of several narrowly delimited taxa to broad, single-species concept (e.g. Richardson 1980; Rix and Woods 1981; Peterson et al. 2010a; Jakab and Molnár 2011; Košťál et al. 2013). Nevertheless, several narrowly delimited taxa are still used in recent literature, mainly in national floras and accounts (Hrouda 2010; Jakab and Molnár 2011; Košťál et al. 2013; Lauber et al. 2018). The notable “narrow” taxa include *G. bohemica* (Zauschn.) Schult. & Schult.f., *G. saxatilis* (Mert. & W.D.J.Koch) Schult. & Schult.f., *G. szovitsii* (Láng) Besser, *G. velenovskyana* Pascher, *G. aleppoana* Pascher, *G. callieri* Pascher, *G. busambarensis* (Tineo) Parl. and *G. bohemica* var. *stenochlamydea* Borbás (Zauschner 1776; Mertens and Koch 1826; Láng 1827; Schultes and Schultes 1829; Borbás 1900; Pascher 1904, 1906; Terracciano 1906; Pascher 1907; Stroh 1936; reviewed by Rix and Woods 1981).

Previous studies of genus *Gagea* indicated poor efficiency of sexual processes, occurrence of proterandry and a strong connection of ploidy level and ability of seed set production (Měsíček and Hrouda 1974; Gargano et al. 2007; Gargano and Peruzzi 2021). Generally, anorthoploids such as triploid *G. lojaconoi* Peruzzi and *G. granatellii*, Parl. and heptaploid *G. fragifera* (Vill.) Ehr.Bayer & G.López showed higher rates of pollen malformation and lower seed set production than the orthoploids such as tetraploid *G. peruzzii* J.-M.Tison and hexaploid *G. lutea* Ker Gawl. (Gargano and Peruzzi 2021). The same pattern was also expected in pentaploid *G. bohemica* from Central Europe which populations usually lack seed sets (Hrouda 1989a; Němec et al. 2017). Němec (1923) made several attempts at controlled fertilisation of Czech and Moravian plants and the only result was the observation that the ovary degenerated after fertilization. In addition to this observation, Měsíček and Hrouda (1974) reported highly irregular meiosis in pentaploid plants of the Bohemian origin. On the other hand, Vardar et al. (2012) reported good ovule development and *Euphorbia dulcis*-type embryo sac in plants of unknown ploidy level from Turkey.

In this study we aimed to 1) infer composition and distribution of cytotypes in *G. bohemica* s. lat. outside the Mediterranean region, 2) investigate cytotype population structure, 3) test whether there are any geographically based differences in genome size that might indicate the evolutionary history of the populations and 4) explore the relationship between the ploidy level and pollen stainability.

## Material and methods

### Plant material

In total, we sampled 106 populations of *G. bohemica* s. lat. covering almost the whole range of distribution outside the Mediterranean (Austria, Bulgaria, Czech Republic, Germany, Hungary, Romania, Serbia, Switzerland and Ukraine) and northern Greece. Field work took place between 2013 and 2023. For the purpose of flow cytometry analyses, we usually took one or two tepals per individual. If there were no flowering individuals in the population, we collected one ground leaf per individual. In several samples, both tepal and leaf were collected in order to infer the possible differences in results from different plant tissue (i.e., due to the presence of higher concentrations of secondary metabolites in one of the tissues). We usually collected 5–10 individuals per population depending on its size (total of 1145 individuals, mean 11 per population, min 1, max 55; Online resource 1). To minimalize resampling of clones, we sampled not obviously connected bulb clusters at least 0.5 m distant from each other. Plants were either kept alive in a refrigerator and analysed as soon as possible or stored in silica-gel until analyses. Several bulbs from selected populations were transferred to cultivation for chromosome counting. For the analysis of pollen stainability, we selected 43 populations from Bulgaria, Czech Republic, Germany, Hungary, Romania and Serbia, a subset of abovementioned populations. Sampled individuals in these populations were the same as for flow cytometry. Two anthers before dehiscence from one individual were usually collected and preserved in a plastic tube with Carnoy’s fixative (6 parts of ethanol : 3 parts of chloroform : 1 part of ice-cold acetic acid). Anthers were usually collected from 5−10 individuals per population depending on its size (total of 289 individuals, mean 7 per population, min 1, max 11; Online resource 1).

### Ploidy level screening

Analysis of DNA-ploidy level (Suda et al. 2006) followed the standard protocol described in Doležel et al. (2007). All sampled individuals (see above) were analysed. About 0.5 cm^2^ of plant tissue was chopped by razor blade in a Petri dish with approximately the same amount of standard in 1 ml of LB01 buffer, pH 7.8 with PVP (polyvinylpyrrolidone; Doležel and Bartoš 2005). *Pisum sativum* ‘Ctirad’ (2C = 9.09 pg) was used as an internal standard. The final solution was filtered through nylon mesh and supplemented either with either 50 μl DAPI (4’,6-diamidino-2-phenylindole) or 50 μl PI (propidium iodide). In the case of DAPI staining, the analysis was performed using a Partec ML flow cytometer (Partec GmbH., Münster, Germany, equipped with UV led diode) and the relative fluorescence of at least 3000 particles was recorded. In the case of PI staining, the analysis was performed using a BD Accuri C6 flow cytometer (BD Biosciences, San Jose, USA) equipped with Accuri blue laser (488 nm, 20 mW, BD Biosciences, San Jose, USA) and at least 3000 (rarely 2000) particles were recorded. Samples from several populations (Online resource 1) were analysed using both flow cytometers to allow comparison between the obtained values from respective machine. The ploidy level of each sample was determined on a linear scale of graphical output by the position of its 2C peak relative to the 2C peak of the internal standard.

### Absolute genome size estimation

The absolute DNA content (absolute genome size, AGS, expressed as 2C), i.e. calculation of genome size in absolute units of picograms according to recommendations given in Doležel et al. (2007), of 46 individuals covering the entire study area (Online resource 1) was determined using a Partec PAS flow cytometer (Partec GmbH., Münster, Germany), equipped with a diode-pumped solid-state green laser (532 nm, 100 mW, Cobolt Samba; Cobolt AB, Stockholm, Sweden). Sample preparation followed the protocol described in full in Kobrlová and Hroneš (2019). Again, *P. sativum* ‘Ctirad’ (2C = 9.09 pg) was used as an internal standard. The fluorescence intensity of at least 5000 particles was recorded. Histograms with coefficients of variation (CVs) of the G0/G1 peaks of both the sample and the standard of less than 5% were accepted. Each individual was analysed at least three times and the average value was used for the calculation of the AGS. The within-measurement variation of one sample was calculated as ((maximum value – minimum value)/average value of all three measurements)*100. If the absolute differences of the three measurements exceeded the 2% threshold, then the most outlying measurement was discarded, and a new measurement was performed. The monoploid genome size, 1Cx value (Greilhuber et al. 2005), was calculated by dividing the 2C value by the ploidy derived from the chromosome counts.

### Chromosome counting

Flow-cytometry results were calibrated by chromosome counts (20 individuals; Online resource 1). Actively growing root tips were collected at the beginning of leaf growth usually during November. Root tips were cleaned of soil residue in distilled water, pre-treated in 8-hydroxyquinoline for 3 hours in the refrigerator and another 3 hours at room temperature in the dark, fixed in 96% ethanol : cold acetic acid (3:1) at least overnight and after that shortly macerated (ca 1 minute) in a solution of 35% hydrochloric acid : 96% ethanol (1:1). Squashes were made in a drop of Fe-acetocarmine and observed under an Olympus BX60 microscope (Olympus, Tokyo, Japan) equipped with an Olympus DP72 camera (Olympus, Tokyo, Japan).

### Pollen stainability

Assessment of pollen stainability is a commonly used marker for evaluating the degree of fertility and meiotic abnormalities of the male gametophyte (Pagliarini 2000; Ramsey and Schemske 2002; Riviero-Guerra 2008; Kolarčik et al. 2015). Counting of stained pollen has also been successfully used to characterize polyploid taxa in genus *Gagea* (Měsíček and Hrouda 1974; Pfeiffer et al 2013; Gargano and Peruzzi 2021). Pollen stainability was tested using a modified Alexander differential stain according to Peterson et al. (2010b) on 289 individuals. One non-dehiscent anther was removed from Carnoy’s fixative, cut by razor blade and the pollen grains were slightly spread on a slide. The rest of fixative was carefully dried out and the plant debris was also carefully removed. One or two drops of the staining solution were applied to the pollen. After 1−2 minutes of staining, the slide was briefly heated over an alcohol burner. A coverslip area of 20 × 20 mm was used for pollen examination under an Olympus CX31 microscope at 200× magnification. All pollen grains under the coverslip were counted using this microscope after the recording the position of a particular field to avoid repeated counts. After counting, photographs of two fields were taken for eventual future data control at 100× magnification using an Olympus BX60 microscope equipped with the QuickPHOTO CAMERA 3.0 program.

### Data analysis

Data were analysed in NCSS 9 (Hintze 2013). The normality of the obtained values was tested by Kolmogorov-Smirnov test. To further test the geographic structure of AGS and pollen stainability, we grouped the populations into three artificial groups based on predominant occurrence of tetraploids (Fig. 2B). Populations from Bulgaria, Greece, southeastern Hungary, Romania and Serbia were grouped together into an eastern group of populations (further abbreviated EE), populations from Austria, Czechia, northwestern parts of Hungary and Slovakia were grouped into a central group of populations (abbreviated CE), and populations from Germany and Switzerland were clustered into a western group of populations (abbreviated WE). Differences in AGS, 1Cx and pollen stainability between the ploidy levels and geographical groups were tested using either ANOVA or Kruskal Wallis test with subsequent use of Tukey HSD test. The maps were created in QGIS 3.28.

## Results

### Karyology and ploidy level screening

Two ploidy levels were discovered by flow cytometry, corresponding to DNA-tetraploids and DNA-pentaploids. Chromosome counts further confirmed these two cytotypes as tetraploid (2n=48; Fig. 3A) and pentaploid (2n=60; Fig. 3B). The results obtained on the basis of different tissues (tepals and leaves) were largely identical..

**Fig. 3.**
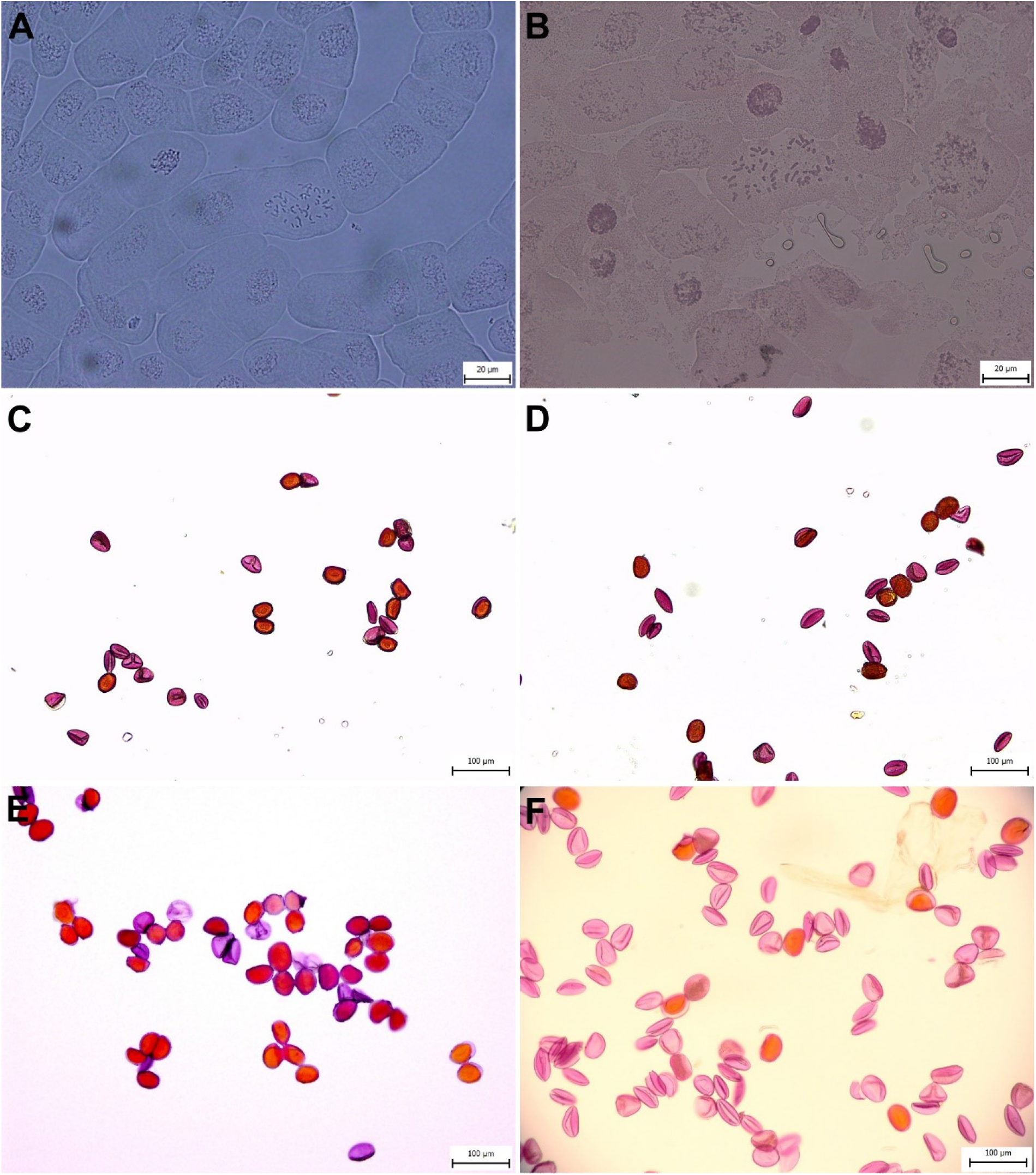
Mitotic metaphase chromosomes and pollen grains of *Gagea bohemica*. Stainable pollen grains are reddish-orange, aborted pollen is pale violet. A – 2n = 4x = 48 (population BOH89). B – 2n = 5x = 60 (population BOH46). C – tetraploid from EE lineage (population BOH18). D – pentaploid from EE lineage (BOH20). E – pentaploid from CE lineage (population BOH25). F – tetraploid from WE lineage (population BOH63).

Tetraploids and pentaploids differ in spatial distribution (Fig. 2B) and occur both in pure (89.6 % of all analysed populations) and mixed-ploidy (10.4 %) populations. Pure tetraploid populations (28.3 %) were found in Germany and Switzerland in the west, and in the Black Sea area (Ukraine, Romania), northern Balkan (Bulgaria, Greece, Serbia) and southeastern Pannonian basin (Hungary) in the southeast of the study area. Two geographically isolated tetraploid populations were discovered in central Moravia (Czech Republic). Pure pentaploid populations (61.3 %) were predominantly concentrated between the two main areas of tetraploid occurrence, in the Bohemian Massif and the Pannonian basin in Central Europe. Two isolated pentaploid populations were found in Switzerland and southern Serbia. Mixed-ploidy populations were mainly present in areas with tetraploid dominance (Germany, southeastern Hungary, south Serbia and Bulgaria).

### Absolute genome size (AGS) estimation

The quality of genome size measurements was generally good. Variation coefficients of standard and sample peaks varied from 1.21 to 4.97 (mean 3.3, SD ± 0.63) and from 1.42 to 4.6 (mean 3.2, SD ± 0.62), respectively. Absolute differences of the three measurements of one sample varied from 0.11% to 1.97% (mean 0.78%, SD ± 0.43).

The AGS (2C) of tetraploids and pentaploids varied from 13.27 pg to 14.1 pg (mean 13.67 pg, SD ± 0.31) and from 15.93 pg to 17.17 pg (mean 16.58 pg, SD ± 0.33), respectively. Significant geographical structure in the AGS was detected between EE, CE and WE population groups (ANOVA, F = 620.06, DF = 5, p < 0.001). Tukey HSD comparison test showed differences in AGS between EE and WE populations (regardless of ploidy), tetraploids from CE populations group differed from EE ones but not WE ones, and pentaploids from the CE populations group differed from WE ones but not EE ones (Fig. 4A, Online resource 2). The monoploid genome size (1Cx) of analysed samples varied from 3.32 pg to 3.53 pg (mean 3.42 pg, SD ± 0.08) in tetraploids and from 3.19 pg to 3.43 pg (mean 3.32 pg, SD ± 0.07) in pentaploids (Fig. 4B) indicating down-sizing in genome size (ANOVA, F = 44.27, DF = 5, p < 0.001). Tukey HSD comparison test showed differences in 1Cx between tetraploids and pentaploids in all three geographic groups (Online resource 2).

**Fig. 4.**
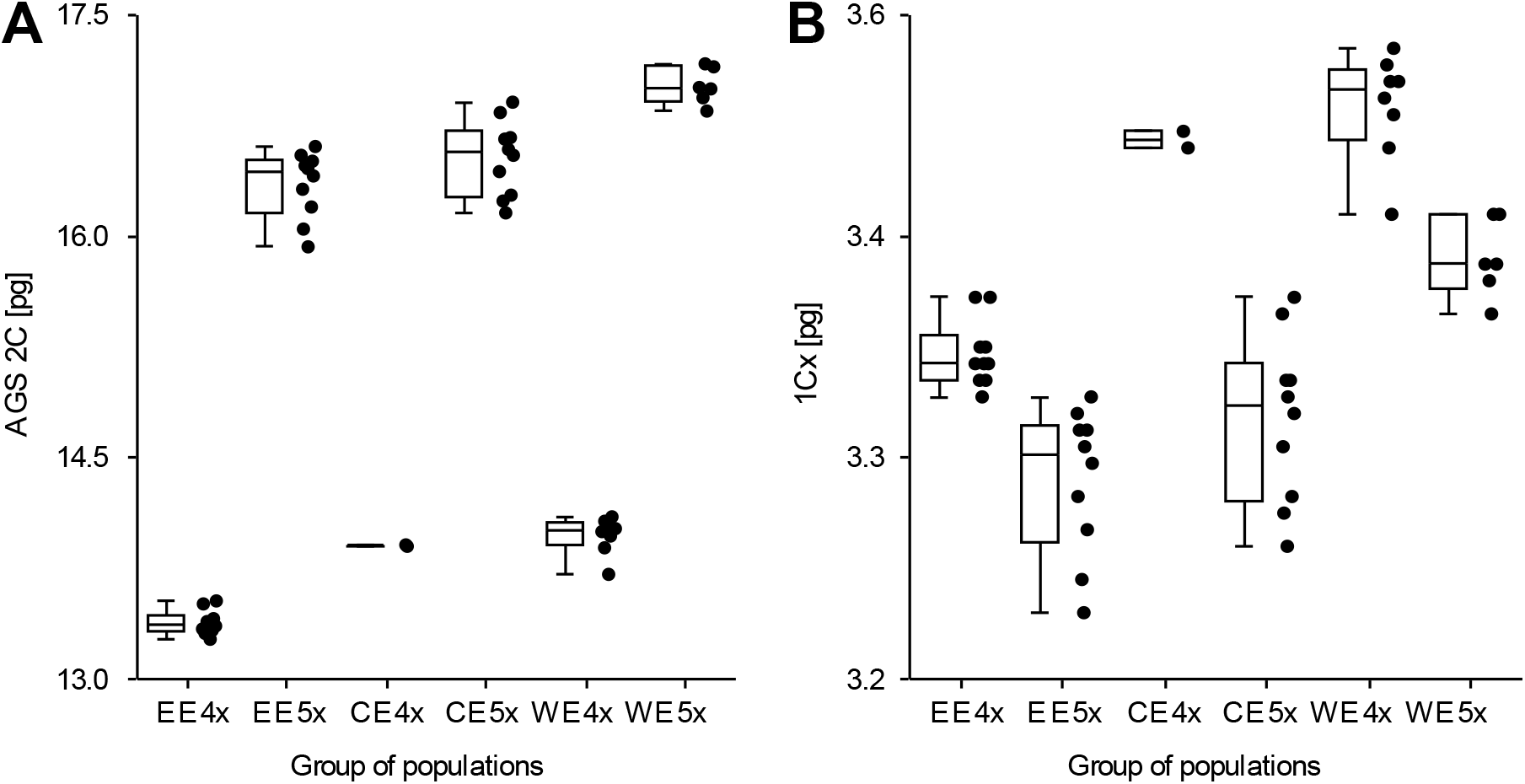
Combined box- and dotplots of established genome sizes in tetraploid and pentaploid *Gagea bohemica* s. lat. sorted according to geography (see chapter Data Analysis for explanation). A – Absolute genome size (2C) values. B – Monoploid genome size (1Cx) values.

### Pollen stainability

In general, tetraploids had on average lower pollen stainability (mean 44.92 %, SD ± 23.33) than pentaploids (mean 72.32 %, SD ± 22.55; Fig. 5A). Pollen grains which were considered aborted using differential staining were generally smaller and deformed. Tetraploids differed in their stainability when divided into geographical groups (χ^2^ = 49.07, p < 0.001). The CE and WE groups had much lower stainability than tetraploids from the EE group of populations. Pentaploids also differed in their pollen stainability (χ^2^ = 57.52, p < 0.001) and all three geographical groups had different stainability from each other with CE group having the highest values and WE group considerably the lowest values (Fig. 5B). Individuals from mixed-ploidy populations regardless of ploidy had either very low (WE group: 4x – mean 29.75 %, SD ± 19.61; 5x – mean 18.89 %, SD ± 11.63) or considerably higher (EE group: 4x – mean 50.35 %, SD ± 14.03; 5x – mean 81.08 %, SD ± 6.72) stainability (Online resource 3).

**Fig. 5.**
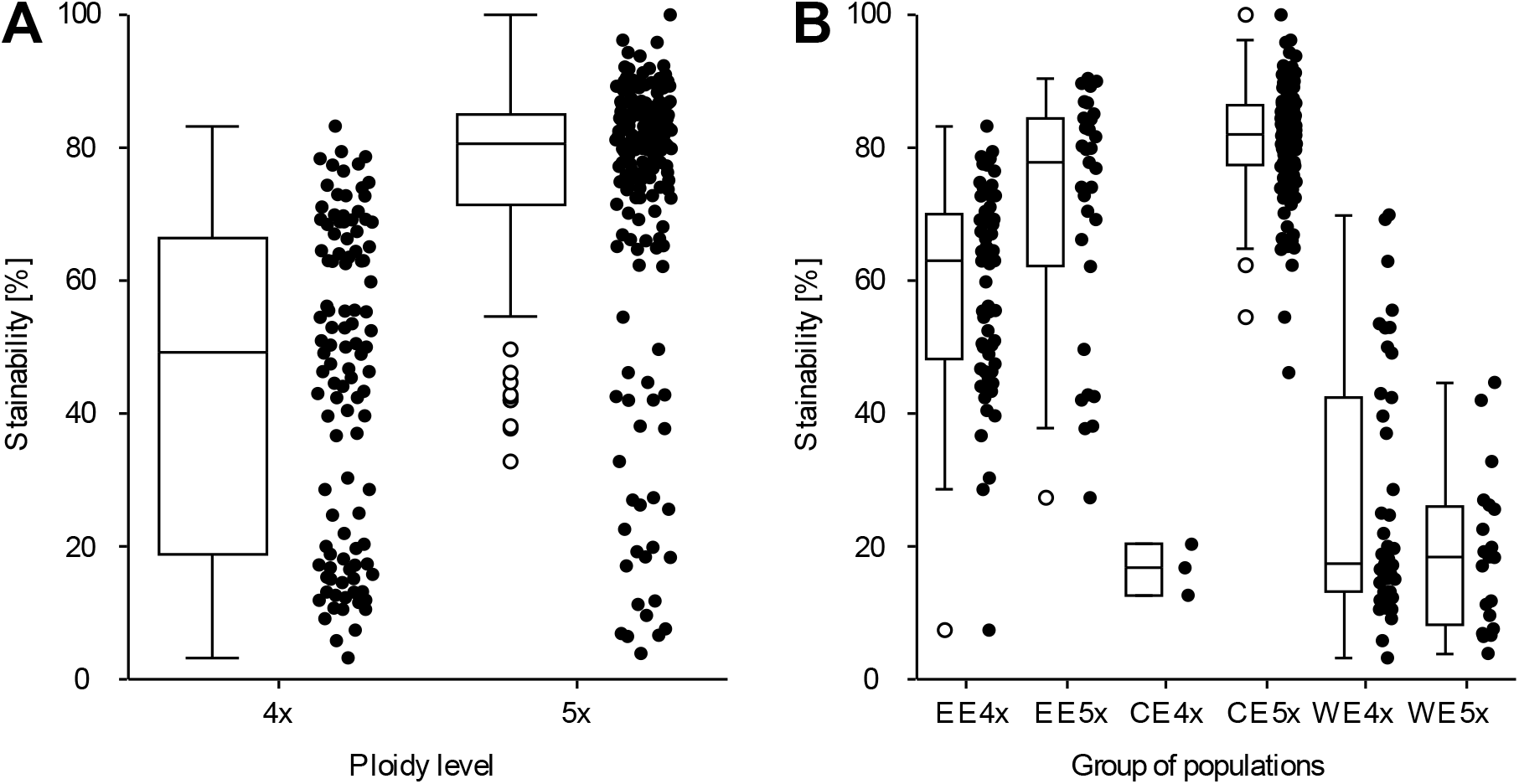
Combined box- and dotplots of proportion of pollen stainability in studied *Gagea bohemica* individuals. A – Comparison of stainability of both cytotypes disregarding their geographic origin. B – Stainability of individuals sorted according to ploidy and geography (see chapter Data Analysis for explanation).

## Discussion

### Cytogeography

*Gagea bohemica* s. lat. is karyologically complex with five reported cytotypes ranging from diploids (2n=24) to hexaploids (2n=72; Fig. 2A). So far, its cytogeography has only been studied on a small scale, mostly in the Mediterranean area of Europe (Peruzzi 2003, 2008) and it is difficult to infer any general pattern from these data. Our analysis of 1145 individuals from 106 populations is, therefore, the first attempt to infer the cytogeography of G. *bohemica* s. lat. on a large scale. In the northern parts of its distribution area, *G. bohemica* s. lat. shows a complex cytogeographic pattern with the occurrence of only two cytotypes – tetraploids and pentaploids. Considering the high karyological diversity of the genus *Gagea* in general (reviewed by Peruzzi 2012) and *G. bohemica* s. lat. in particular, this result is somewhat surprising. Nevertheless, its low karyological diversity is in line with previously reported chromosome counts from this region which include only pentaploids from Czech Republic and Slovakia, and tetraploids from Macedonia (Měsíček and Hrouda 1974; Murín and Májovský 1983; Sopova et al. 1984a, 1984b). Our study adds new chromosome counts for *G. bohemica* s. lat. from Bulgaria, Germany, Romania and Serbia.

We revealed two geographic areas with dominance of tetraploids outside the Mediterranean, which can be referred to as western (WE) and eastern (EE) ones. These two areas coincide with the reported presence of diploids and triploids in the western (France and Corse) and the eastern Mediterranean (Israel and Turkey; Fig. 2A; Contandriopoulos 1962; Heyn and Dafni 1977; Guerlesquin 1985; Tison 1996; Özhatay 2002). Given the difference between the western and eastern tetraploid lineages in AGS, it is possible that they represent two independent lineages arising from different refugia in the Last Glacial Maximum. Similar pattern with two lineages differing in their genome size with presumable origin from two refugia was discovered in *Allium oleraceum* L. (Duchoslav et al. 2020). However, to further confirm our hypothesis the population with reported diploid and triploid counts need to be studied first to confirm their taxonomic identity and the presence of such counts in their populations. *Gagea bohemica* s. lat. populations also differ in their ecology in these two areas, which may further confirm their independent origin. Populations in Germany (Fig. 1E), France and Switzerland occupy sandy and/or shallow porphyritic, melaphyric or weathered shale soils with frequent occurrence of (sub)oceanic species (Korneck 1975; Landolt et al. 2010). On the other hand, tetraploids in southeastern Europe frequently inhabit nutrient-poor alkaline grasslands covered by continental species (Fig 1F; Jakab and Molnár 2011). In addition, at least some of *G. bohemica* populations from Bulgaria and Serbia sampled in this study originated from alkaline rock outcrops. This could indicate that EE lineage occupies habitats with a higher pH more often than WE lineage. Purely pentaploid populations in CE group of populations occupy an intermediate geographical position between the two areas with predominantly tetraploid lineages. These populations grow in habitats similar to the ones of WE lineage (see Černý et al. 2011). They also possess slightly higher AGS values in average then EE group pentaploids and considerably lower AGS values than WE group pentaploids which puts them into intermediate position when compared to pentaploids from regions of tetraploid dominance (i.e., EE and WE groups). There are several possible scenarios of their origin. The CE populations may be descendants of EE populations which migrated further northwest. Their slightly higher AGS can be explained by the accumulation of transposable elements along the way. Another possible explanation of their origin which, may also explain their AGS values, is that they arose from hybridization between EE and WE lineages.

### Mixed-ploidy populations

Mixed-ploidy populations of one species provide an interesting insight into cytotype interactions, evolution and ecology (Kolář et al. 2017). The mixed-ploidy populations in *G. bohemica* s. lat. have not been reported before our study. Because of only a few flow cytometric studies on the genus *Gagea*, this phenomenon has remained unexplored. Even more curious is the coexistence of tetraploid and pentaploid and the absence of hexaploid cytotypes in mixed-ploidy populations. Despite considerable effort, we did not find any other cytotype in *G. bohemica* s. lat. populations in the study area. A similar pattern of mixed-ploidy populations consisting of tetraploids and pentaploids is very unusual in plants in general, but it was also found in other monocot geophytes, such as *Allium oleraceum* (Duchoslav et al. 2010). It is rather difficult to explain the observed pattern of cytotype coexistence. Due to the presence of genome down-sizing in pentaploids when compared to tetraploids, i.e. the lower monoploid genome size values of pentaploids when compared to tetraploids, the origin of mixed-ploidy populations is hard to establish. However, we expect that mixed-ploidy populations are a result of primary contact (i.e. on-site) between cytotypes rather than secondary one (i.e. result of migration after allopatric origin). The current state hints on presence (at least historical) of other cytotypes in the surveyed populations. Rare hexaploid formation in a tetraploid population via fusion of reduced and unreduced gametes from a tetraploid and subsequent backcrossing of a reduced gamete from hexaploid with a reduced gamete of tetraploid can explain the observed pattern. An important factor that contributes to the long-term survival and coexistence of both cytotypes in mixed-ploidy populations is the ability of *G. bohemica* s. lat. to propagate vegetatively through bulbils (Hrouda 2010; Elias et al. 2018). The conspicuous absence of mixed-ploidy populations in CE group was previously suggested to be a result of the extinction of pollinators in last glacial period, which led to the extinction of euploid cytotypes and subsequent survival of anorthoploid, vegetatively reproducing cytotypes only (Hrouda 1989b).

### Ploidy level vs. reproduction and implication for population management and conservation

Ploidy level plays an important role in the ability of individual to successfully reproduce and to produce viable offspring (e.g., Thompson et al. 2008; Duchoslav and Staňková 2015; Sattler et al. 2016). According to the available literature, pentaploids of *G. bohemica* s. lat. heavily depend on vegetative propagation via bulbils and reports of their seed set are very rare (Němec 1923; Slater 1990; Hrouda 1989a, 2010). On the contrary, tetraploids can produce at least occasionally a small amount of seeds (Caparelli et al. 2006; Gargano et al. 2007; Peterson et al. 2010a; Lambelet and Détraz-Méroz 2018). The frequent seed set occurrence was also reported from southeastern Hungarian populations (Jakab and Molnár 2011) where we revealed that tetraploid individuals are very frequent. Thus, the ability to produce seeds in *G. bohemica* s. lat. is clearly affected by ploidy level rather than the lack of pollinators as stated by Uphof (1959). On the other hand, we observed quite unusual pattern of pollen stainability with tetraploids having reduced stainability while pentaploids having a much higher proportion of stainable pollen. The cause of reduced stainability in even ploidy can be explained in some populations by their presumed low genetic variability resulting from low number of individuals as in populations of tetraploids from Moravia (e.g., Paschke et al. 2002), but in others a simple mechanistic explanation is lacking. To our knowledge, a similar pattern of reduced pollen stainability/viability in even ploidy cytotypes and high stainability in odd ploidy cytotypes has been seldom reported but is known in *Ornithogalum* (Raamsdonk 1985). On the other hand, it is in stark contrast to the observations made by Pfeiffer et al. (2013) on *G. lutea* (generally hexaploid), *G. pratensis* (generally pentaploid) an their hexa- and heptaploid hybrids, where *G. pratensis* and the heptaploid hybrids had lower pollen viability compared to hexaploid hybrid and *G. lutea*. A similar situation was also reported by Měsíček and Hrouda (1974) for tetraploid and pentaploid plants from the *G. pratensis* group. Latter authors also evaluated stainability in one accession of pentaploid *G. bohemica* and reported a very similar result to ours (stainability of approximately 80 %). Even more interesting is the different pattern of stainability in the eastern and the western tetraploid lineages. Individuals from the eastern tetraploid lineage have considerably higher stainability than the western ones, and the proportion of their pollen stainability almost reaches the pollen stainability of pentaploids, regardless of their occurrence in cytotype pure or mixed-ploidy populations. On the contrary, western tetraploids as well as their sympatric pentaploids have reduced stainability. The low stainability of these populations cannot be explained by sampling bias (e.g., sampling of frost-damaged anthers) as our samples were collected in several different years. One possible explanation is the very low genetic diversity of these populations. The other is the presence of aberrant meiosis in pentaploids which create gametes of different ploidies (e.g., 2x, 3x, 4x). Diploid gametes from pentaploids may merge with normal gametes (2x) of tetraploids. The resulting tetraploid offspring may consequently suffer from aberrant meiosis and reduced genetic variability which is further pronounced in their very low pollen stainability.

Vegetative dispersion and the inability to produce seeds in pentaploids is pronounced by low genetic diversity in their populations (Peterson and Peterson 1999). Knowledge of ploidy level is therefore extremely important for the successful nature conservation management application when increased seed set is considered a desirable outcome. This can be illustrated on the study of Elias et al. (2018) who observed very low seed set despite the application of two management types. We resolved one of the three populations they studied (Mücheln; BOH64) as mixed-ploidy with a high proportion of pentaploids and discovered several other mixed-ploidy populations in the same region. It is therefore not surprising that the seed set reported by Elias et al. (2018) on population level was low.

### Implications for taxonomy

Several contrasting taxonomic concepts has been proposed for *G. bohemica* s. lat. ranging from the recognition of several “narrow” taxa to a single polymorphic species (Pascher 1904, 1906, 1907; Terracciano 1906; Richardson 1980; Rix and Woods 1981; Peterson et al. 2010a). The studied area covers regions from where several “narrow” taxa of *G. bohemica* s. lat. were described in the past, including the type area of *G. bohemica* in Central Bohemia (Kirschner et al. 2007). The plants from Bohemia together with the majority of plants from the northern parts of the Pannonian basin formed purely pentaploid populations with a fairly homogeneous AGS. Another “narrow” taxon, *G. szovitsii*, was described from Ukraine but later reported also from Romania and south-eastern Hungary (Richardson 1980; Jakab and Molnár 2011). Plants from these areas forms purely tetraploid, purely pentaploid and mixed-ploidy populations with the lowest AGS values within the analysed data set and with high pollen stainability. Populations of *G. bohemica* s. lat. from the Plovdiv region of Bulgaria were described as *G. velenovskiana* (Pascher 1906). We had the opportunity to study the population near Vinogradec village not far from Plovdiv which is tetraploid. *Gagea saxatilis* was described from Germany and the name was later extended to western European (from Portugal to Germany), central European (Czechia) and southeastern European (Bulgaria, Macedonia, Greece) populations (Richardson 1980; Assyov et al. 2006; Hrouda 2010; Jäger 2017; Lauber et al. 2018). The prevailing ploidy level of the populations that are assigned to this taxon is tetraploid with frequent occurrence of mixed-ploidy populations. However, the AGS of all populations from Germany and Switzerland were significantly higher than that of central and eastern European populations suggesting non-homogeneity of some taxonomic concepts considering *G. saxatilis*. Similarly high AGS of plants from two isolated tetraploid populations from Central Moravia in Czechia suggests their affinity to the western European ones. This is also in line with the assignment of these populations to “western type” (as *G. bohemica* subsp. *saxatilis*) based on their morphology (Hrouda 2010; Horák et al. 2017). When confronted with data from Peterson et al. (2010) which partly cover the same populations of *G. bohemica* s. lat. as this study, the cytogeographic pattern does not correspond to observed *trn*L-*trn*F IGS haplotypes. However, their study does not cover the populations in southeastern Europe which represent the EE group of populations and may poses an undiscovered haplotype variation. Discrepancy of our data obtained by flow cytometry and chloroplast sequencing given by Peterson et al. (2010) encourage more precise in-depth genome sequencing on wider sample set to resolve the evolution of *G. bohemica* s. lat.

The overall pattern of cytotype distribution and AGS variation may support the distinction of three geographically delineated groups/lineages (i.e., western predominantly tetraploid with high AGS, central pentaploid with AGS values similar to eastern ones and eastern tetraploid/pentaploid with low AGS) as separate taxa, probably at a lower taxonomic level than species. Alternatively, only two groups may be recognised based on similarity of AGS values and pattern of pollen stainability (western one from Germany and Switzerland including two tetraploid populations from Moravia and eastern one from southeastern Europe including pentaploid populations from Central Europe). Given the complicated nomenclature, the high number of described taxa (reviewed by Rix and Woods 1981) and the conflicting taxonomy of the group in different parts of its range (see Online resource 1), we refrain from a proposing a taxonomic concept here and we emphasize that these abovementioned hypotheses need to be further tested using morphology and DNA-based analyses.

## Supporting information

Online resource 1

Online resource 2

Online resource 3

## Acknowledgement

We are indebted to Lucie Kobrlová for help with field work and flow cytometry analyses, Martin Duchoslav for discussion on the earlier versions of the manuscript and Lubomír Hrouda for providing his unpublished manuscript. Two anonymous reviewers are thanked for their comments and suggestions to the earlier versions of the manuscript. We are also very grateful for help with sample collections and/or fieldwork by following colleagues: Pavol Eliáš jun. (Nitra, Slovakia), Gabriel Gigea (Limanu, Romania), Vasilis Ioannidis (Kilkis, Greece), Judit Sallainé Kapocsi (Körös-Maros NP, Szarvas, Hungary), Ivan Kostadinov (Sliven, Bulgaria), Jaroslav Košťál (Nitra, Slovakia), Ivan Moysiyenko (Kherson, Ukraine), Zdeněk Musil (Blansko, Czech Republic), Radomír Němec (Znojmo, Czech Republic), Mirjana and Miloš Petrović (Kruševac, Serbia), András Schmotzer (Eger, Hungary), Tomáš Tichý (Karlštejn, Czech Republic) and Vojtěch Žíla (Strakonice, Czech Republic). Work of BT, DH and MH was supported by Internal grant agency of Palacký University grant IGA PrF-2023-001, DH was further supported by DAAD nr. 57507442. Work of TV was supported by the Ministry of Agriculture of the Czech Republic within the framework of the Long-Term Conception of the Development of the Research Organization Agricultural Research, Ltd. Troubsko.

## Information on electronic supplementary material

### Online resource 1

Overview of sampled localities including numbers of individuals used for each analysis, genome sizes and mean values of stainability and sample/standard ratios from flow cytometry.

### Online resource 2

Results of Tukey HSD multiple comparison of absolute genome sizes (AGS) and monoploid genome sizes (1Cx) between geographical groups and cytotypes.

### Online resource 3

Pattern of pollen stainability in pure and mixed-ploidy populations sorted according to their geographic affiliation.

