## Supplementary material for "Cytogeography of *Gagea bohemica* (Liliaceae) outside the Mediterranean: two ploidy levels, spatial differentiation of cytotypes, and occurrence of mixed-ploidy populations": Online resource 2

**Online resource 2** Results of Tukey HSD multiple comparison of absolute genome sizes (AGS) and monoploid genome sizes (1Cx) between geographical groups and cytotypes.

Absolute genome size:

Alpha=0.050 Error Term=S(A) DF=40 MSE=0.03185525 Critical Value=4.2374

Group	Count	Mean	Different From Groups

CE4x	2	13.905	CE5x, EE4x, EE5x, WE5x

CE5x	10	16.534	CE4x, EE4x, WE4x, WE5x

EE4x	10	13.382	CE4x, CE5x, EE5x, WE4x, WE5x

EE5x	10	16.352	CE4x, EE4x, WE4x, WE5x

WE4x	8	13.975	CE5x, EE4x, EE5x, WE5x

WE5x	6	17.02	CE4x, CE5x, EE4x, EE5x, WE4x

Monoploid genome size:

Alpha=0.050 Error Term=S(A) DF=40 MSE=0.001330833 Critical Value=4.2374

Group	Count	Mean	Different From Groups

EE4x	10	3.346	EE5x, CE4x, WE4x, WE5x

EE5x	10	3.27	EE4x, CE4x, WE4x, WE5x

CE4x	2	3.475	EE4x, EE5x, CE5x

CE5x	10	3.307	CE4x, WE4x, WE5x

WE4x	8	3.495	EE4x, EE5x, CE5x, WE5x

WE5x	6	3.4033	EE4x, EE5x, CE5x, WE4x
