## Supplementary material for "Cytogeography of *Gagea bohemica* (Liliaceae) outside the Mediterranean: two ploidy levels, spatial differentiation of cytotypes, and occurrence of mixed-ploidy populations": Online resource 3

**Online resource 3** Pattern of pollen stainability in pure and mixed-ploidy populations sorted according to their geographic affiliation.

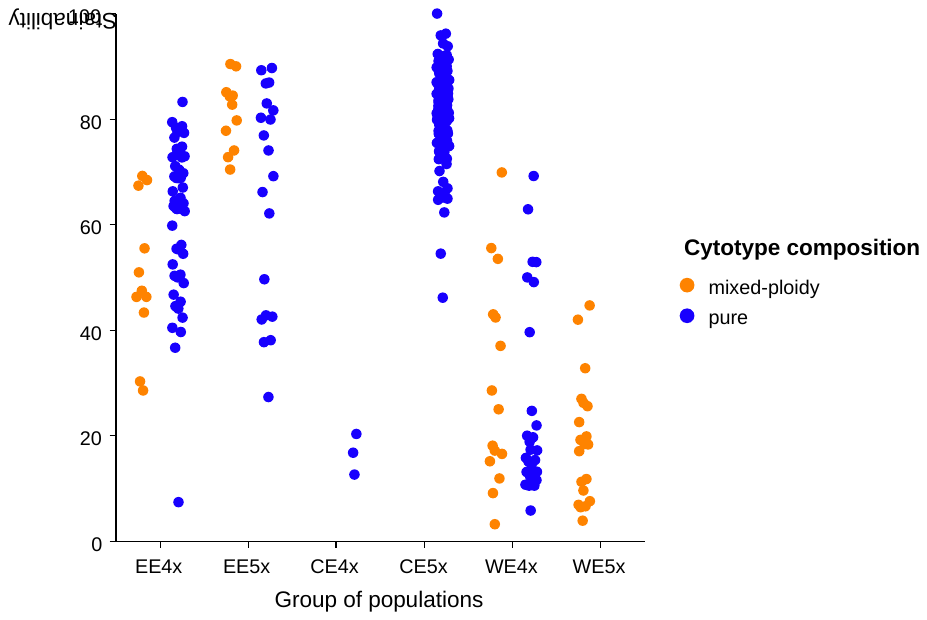
